# Hemoxygenase-1 as a key mediator of acute radiation pneumonitis revealed in a human lung alveolus-on-a-chip

**DOI:** 10.1101/2023.02.13.528339

**Authors:** Queeny Dasgupta, Amanda Jiang, Sean Hall, Robert Mannix, Amy M. Wen, Donald E. Ingber

## Abstract

Exposure to gamma radiation either due to environmental disasters or cancer radiotherapy can result in development of acute radiation syndrome (ARS), characterized by pneumonitis and lung fibrosis. We leveraged a microfluidic organ-on-a-chip lined by human lung alveolar epithelium interfaced with pulmonary endothelium to model acute radiation-induced lung injury *in vitro*. Both lung epithelium and endothelium exhibited DNA damage, cellular hypertrophy, upregulation of inflammatory cytokines, and loss of barrier function within 6 h of radiation exposure, although greater damage was observed in the endothelium. Transcriptomic analysis revealed increased expression of the cytoprotective gene, hemoxygenase-1 (HMOX-1) and gene network analysis identified it as a central mediator of radiation-induced injury. Pharmacological stimulation of HMOX-1 activity also significantly reduced acute radiation-induced lung injury, although it enhanced damage at later times. Thus, this human lung chip offers a new platform to study ARS and these results suggest that HMOX-1 may be mechanistically involved in this injury response.

## INTRODUCTION

Radiation induced lung injury (RILI) can occur in radiation disasters and it is also can be a toxic side effect of radiation therapy, which is administered to ~60% of cancer patients [1]. Radiation injury in lung cells causes DNA damage and accumulation of reactive oxygen species (ROS), which induces an inflammatory response [2, 3]. Sustained inflammation causes alveolar epithelial cell damage accompanied by severe injury to the surrounding endothelium. This inflammatory phase is called radiation-induced pneumonitis (RP) and occurs between 1-3 weeks of exposure [2, 4]. RP is commonly diagnosed based on clinical symptoms, including shortness of breath, dry cough, low-grade fever, chest pain or general malaise. However, no specific diagnostic tests or imaging approaches can definitively correlate findings to clinical symptoms [4]. At present, most patients are given prolonged glucocorticoid therapy using prednisone to reduce inflammation over 4-12 weeks after exposure [5]. Unfortunately, there is limited evidence to support its effectiveness at reducing pneumonitis, although there can be some small benefits with regards to subsequent fibrosis [6]. There is, therefore, a great need for clinically relevant preclinical models that may be used to better understand the molecular basis of RILI as well as for the development of improved diagnostics and radiation countermeasure therapeutics.

Animal models have been employed to understand RILI, but their failure to recapitulate key hallmarks of the human pathophysiology limit their value [7]. For example, mouse RILI models only exhibit low grade pneumonitis and minimal pulmonary fibrosis [8–10]. These studies also typically use animal survival as the primary experimental endpoint [11], instead of incidence and severity of pneumonitis, which is more human-relevant. Currently, non-human primate (NHP) models are considered the gold-standard for radiation injury [12, 13], but their use is limited by short supply, long breeding periods, high costs, and serious ethical concerns [14, 15]. *In vitro* models comprising lung epithelium exposed to radiation provide some insight to the effects of radiation on individual cell types, but do not provide more physiologically relevant information about inflammation responses, destruction of tissue barriers, and the interplay between different cell types during RILI that are observed in the human body [16–18].

In the present study, we leveraged organ-on-a-chip (Organ Chip) microfluidic culture technology to model RILI in the human lung in vitro. Organ Chip models of lung alveolus and airway have been successfully used in the past to model multiple lung diseases, including asthma, chronic obstructive pulmonary disease (COPD), pulmonary thrombosis [19], cystic fibrosis [20], and lung cancer [21], as well as viral infection and evolution of resistance to antiviral therapy [22, 23]. Here, we used a previously described 2-channel Lung Alveolus Chip containing primary human lung alveolar epithelial cells cultured under an air-liquid interface (ALI) interfaced across an extracellular matrix (ECM)-coated porous membrane with a primary human lung microvascular endothelium that lines a channel which is dynamically perfused with culture medium with or without circulating human immune cells [22,24,25]. The engineered alveolar-capillary interface also experiences cyclic mechanical strain to mimic breathing motions. Importantly, we show that when this human Lung Alveolus Chip is exposed to clinically relevant doses of gamma radiation, many of the physiological hallmarks of RILI are observed. This in vitro lung alveolus model also recapitulates the known effects of radiation on induction of hemoxygenase-1 (HMOX-1) as well as the protective effects of a known radiation countermeasure drug (prednisolone) [2]. Moreover, using this model, we show that experimental elevation of HMOX-1 protects the alveolus against radiation injury, immediately after injury, indicating that HMOX-1 is mechanistically involved in the RILI response.

## RESULTS

### Modeling radiation-induced pneumonitis in the Lung Alveolus Chip

The Lung Alveolus Chip is a microfluidic culture device contains two microchannels separated by a porous ECM-coated membrane lined by human primary alveolar epithelial cells on one side and human lung microvascular endothelial cells (HMVEC-L) on the other to mimic the alveolar-capillary interface of human lung (**Fig. 1a**). Immunostaining for ZO-1 in tight junctions and E-cadherin in cell-cell adhesions in alveolar epithelial cells and VE-cadherin in endothelial cell-cell adhesions, along with DAPI staining of nuclei, confirmed the presence of a continuous alveolar-capillary interface with the endothelium lining all four sides of the lower channel beneath the porous membrane with a monolayer of alveolar epithelium above (**Fig. 1a** and **Supplementary Fig. S1a**). The experimental protocol (**Fig. 1b**) involves culturing the cells for 14 days at an ALI in the presence of cyclic mechanical strain to mimic breathing motions in order to ensure optimal differentiation (see **Methods**) before exposure to gamma radiation on Day 15.

**Fig. 1:**
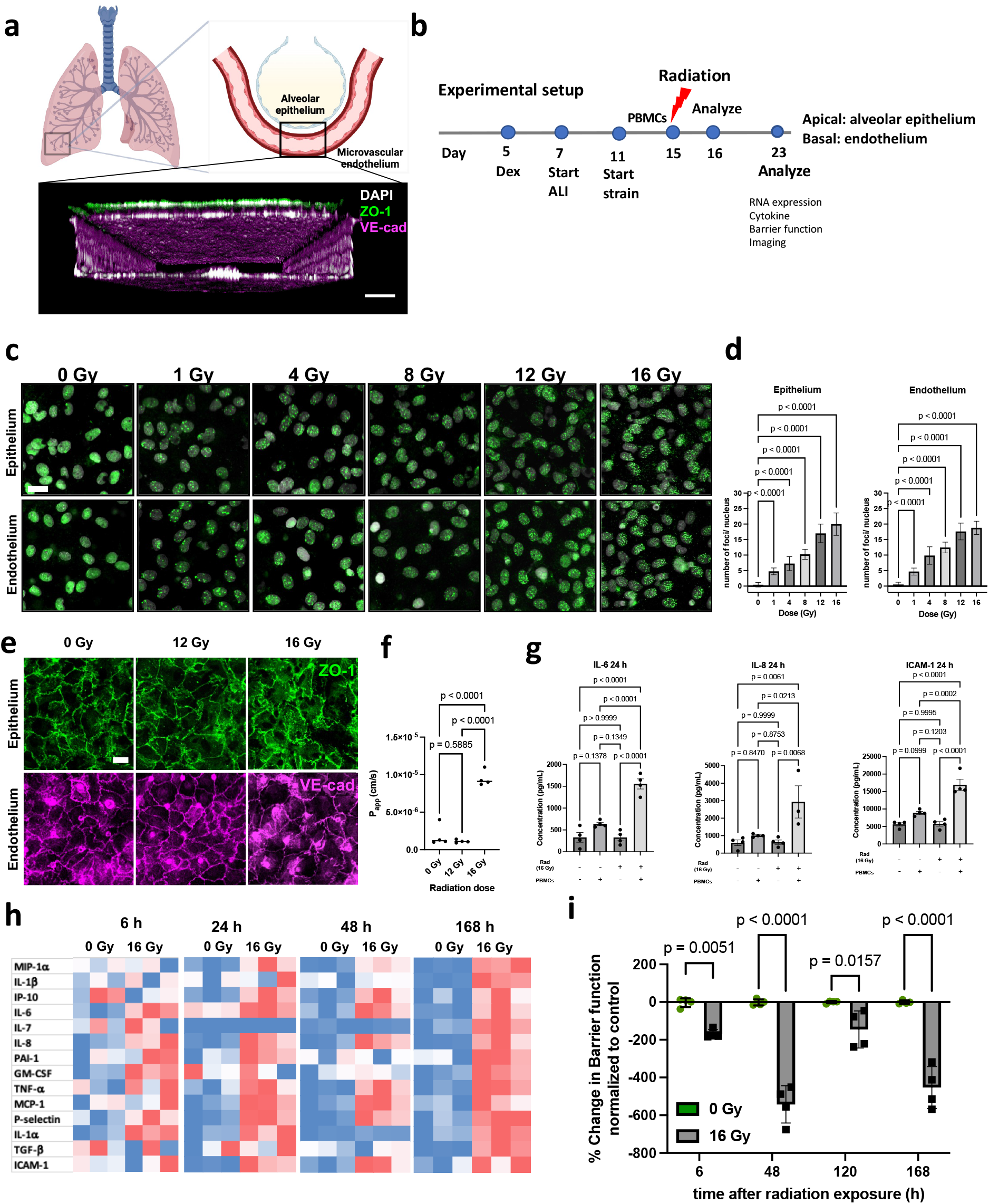
Human lung alveolus chip recapitulates hallmark features of RILI. (a) Schematic of the alveolus-on chip model (created with Biorender.com), showing the confocal z-stack illustrating endothelial tube formation. Scale bar = 100 μm. (b) Line diagram showing experimental plan, Dex, dexamethasone; ALI, air-liquid interface. (c) 53bp1 immunostaining (green) for double-stranded DNA breaks 2 h after radiation. DAPI counterstaining is shown in white. Scale bar = 20 μm. (d) Formation of 53BP1 foci was quantified per cell and showed a dose-dependent increase for both alveolar epithelial and endothelial cells. Data shown are mean +/- S.D. (no. of nuclei per group n=50; one-way ANOVA DF=5, F=525 (e) Immunostaining with ZO-1 (epithelial cells; green) and VE-cadherin junctions (endothelial cells; magenta) post irradiation, showed that junction disruption required a minimum dosage of 16 Gy. Scale bar = 20 μm. (f) Barrier function assay showed a 7-fold increase in the apparent permeability co-efficient (Papp) at 6 h post radiation exposure, but no difference in response to 12 Gy. One-way ANOVA: n= 4, DF= 2, F= 80.6 (g) Representative Comparison of cytokine levels, 24 h post-radiation shows an inflammatory response to 16 Gy radiation, in the presence of PBMCs. One-way ANOVA, n=4, F(IL-6) = 40.05, F(IL-8)= 7.74, F (ICAM-1)= 34.07 (h) Heatmap showing cytokine response to radiation at 6 h, 24 h, 48 h and 7 d post radiation exposure, in the presence of PBMCs (i) % change in barrier integrity normalized to 0 Gy control over 7 days post-radiation exposure, in the presence of PBMCs. 2-way ANOVA, n= 4, DF=3, F= 19.3.

Ionizing radiation induces damage to the cells by generating DNA double-strand breaks (DSBs), leading to hyperphosphorylation of DSB-associated proteins, which produce discrete nuclear foci containing p53-binding protein 1 (53bp1) [24, 25]. When we exposed the Alveolus Chips to increasing doses of radiation from 0 to 16 Gray (Gy), we observed a dose-dependent rise in DNA damage (DSB formation) in both the epithelium and endothelium, as measured by quantifying the number of 53bp1-positive foci per nucleus (**Fig. 1c,d**). Interestingly, there was a significant increase in DSB formation with radiation exposures as low as 1 Gy (**Fig. 1c, d**).

While both the alveolar epithelium and endothelium appeared as regular monolayers composed of cells surrounded by continuous cell-cell junctions in control chips (**Fig. 1a, e** and **Supplementary Fig. S1a**), damage to the tight junctions was only observed when the chips were exposed to the highest radiation dose (16 Gy), but not at the lower exposures that induced DNA damage (**Fig. 1e** versus **Fig. 1d**). This was further corroborated by quantifying tissue barrier function, which showed that the apparent permeability (Papp) of the alveolar-capillary interface increased by ~7 fold after 6 h exposure to 16 Gy radiation (**Fig. 1f**), and this was accompanied by fluid accumulation in the epithelial channel between 5 to 7 days following radiation exposure in ~85% of the radiated chips. Thus, the Alveolus Chip recapitulates the compromise of lung tissue barrier function that is a critical feature of RILI, which emerges about a week after exposure to a similar high dose of radiation in humans.

As inflammation is also a common feature of RILI, we analyzed protein expression levels of multiple proinflammatory cytokines in chips exposed to radiation. However, we could not detect any changes in key proinflammatory cytokines (IL-6, IL-8, TNF-α, GM-CSF) or leukocyte adhesion markers (ICAM-1, E-selectin) even at the highest 16 Gy dose (**Supplementary Fig. S1b**). This observation raised the possibility that the lack of this response was due to the absence of immune cells.

To explore this possibility, we performed similar studies while perfusing the endothelium-lined vascular channel with primary human peripheral blood mononuclear cells (PBMCs) before and during exposure to radiation. Importantly, the presence of PBMCs at the time of radiation exposure caused a sustained and progressive inflammation response in the Alveolus Chip (**Fig. 1g**). We observed upregulation of IL-8, IL-1α, P-selectin, IL-6, PAI-1, MIP-1a, TNF-α, ICAM-1 at 1 day (**Fig. 1g**) and several other cytokines at 1 to 7 days after radiation exposure (**Fig. 1h**). While the inflammatory response was similar at 1 and 2 days, the Alveolus Chips displayed a heightened response to radiation in both expression levels and in the number of affected cytokines at day 7 (**Fig. 1h**). For example, IL-1β, IL-7, GM-CSF and TGF-β were only overexpressed at this later time point. These findings are consistent with the clinical observation that the onset of RP occurs about 1 week after injury [2]. We also observed that barrier permeability (P_app_) was significantly higher in radiated samples when PBMCs were present compared to control (**Fig. 1i**) and this correlated with increased disruption of PECAM1-containing cell-cell junctions in the endothelium (**Supplementary Fig. S1c**). Owing to the importance of PBMCs to generate an inflammation response, all subsequent radiation experiments on the Alveolus Chip were performed in the presence of PBMCs.

Analysis of acute cellular responses to radiation in the presence of PBMCs revealed significant increases in ROS levels within 2 hours (**Fig. 2a)**, as well as hypertrophy of the alveolar epithelial cells at 6 hours (**Fig. 2b,c),**which is a feature commonly seen in alveolar injury in vivo. While cellular hypertrophy was not detected in the endothelium at this early time point, the size of both the epithelial and endothelial cells increased significantly by day 7 (**Fig. 2b,c)**

**Fig. 2:**
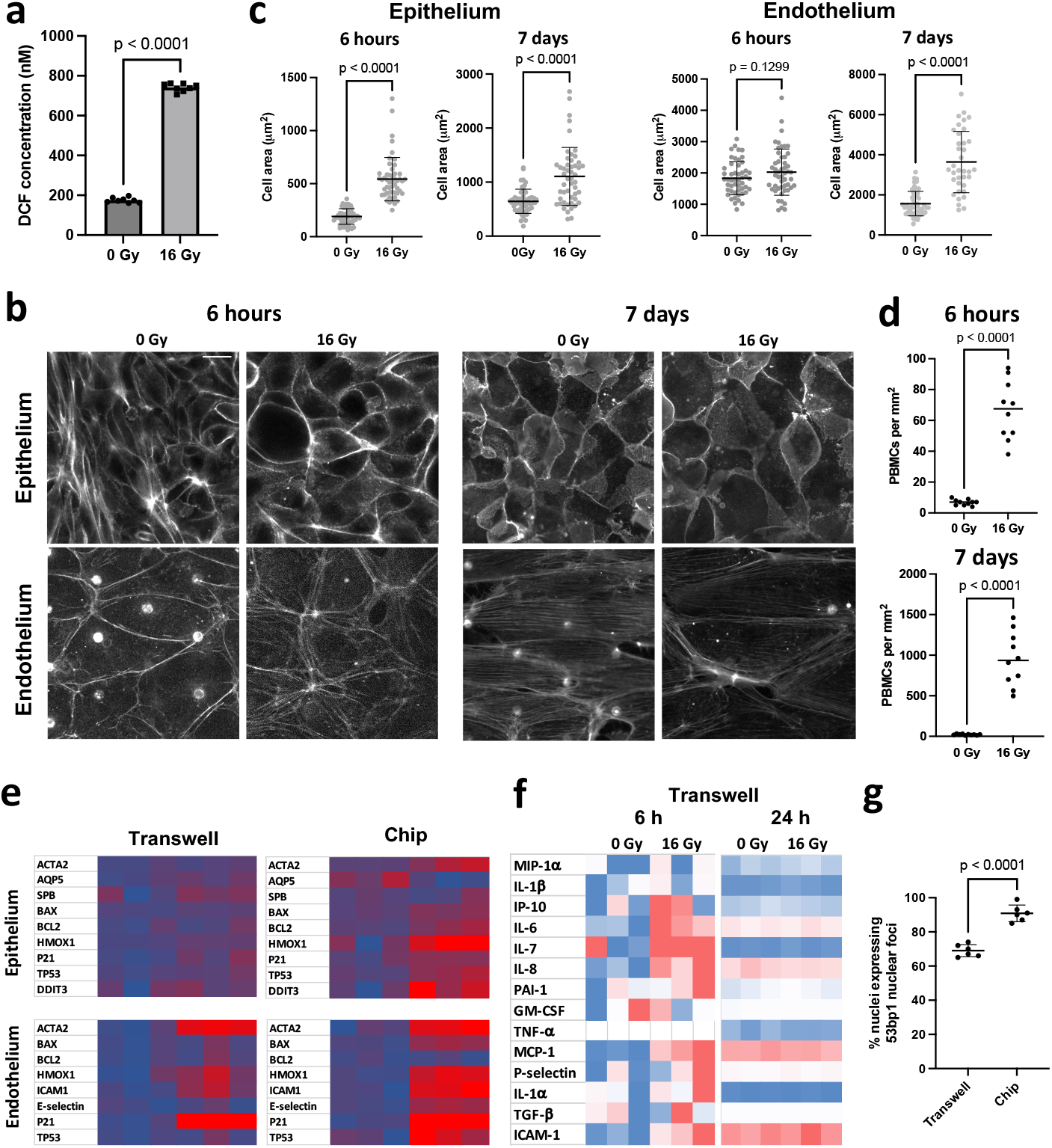
Radiation injury causes ROS, cellular hypertrophy and increased recruitment of PBMCs over time on the alveolus chip. (a) 16 Gy radiation induces increase in the ROS levels, 2 h after radiation injury, n= 4 chips (b) IF images showing the effect of radiation on cellular hypertrophy in the epithelium and endothelium 6 h and 7 d post radiation exposure, Scale bar = 20 μm. Corresponding quantification of cellular area in the epithelium and endothelium (c). (d) Quantification of PBMC recruitment to the vascular endothelium at 6 h and 7 d post-radiation exposure. (e) Heatmaps comparing gene expression profiles in Transwells and chip. The chip shows susceptibility to radiation that is evidenced by the overexpression of several markers. (f) Heatmap showing cytokine profiles of transwells at 6 h and 24 h post-radiation. At 6 h, the transwells exhibit an upregulation of IL-6, IL-8 and ICAM-1 but at 24 h, no effect of radiation exposure is observed. (g) Quantification of % nuclei expressing nuclear foci in response to radiation in transwells vs chips. Two-tailed Student’s t-test with Welch’s correction.

We then investigated how radiation exposure influences PMBC recruitment over time. When immune cells were perfused through the vascular channels of the Lung Alveolus Chip and exposed to 16 Gy radiation, 10 times more PBMCs bound to the endothelium compared to control non-radiated chips when analyzed 6 hours later, and this recruitment increased by almost 5-fold more (levels in radiated chips ~46 times higher than control) by 7 days after radiation (**Fig. 2d** and **Supplementary Fig. S1d**).

Importantly, when similar studies were carried out by adding PBMCs to irradiated static Transwell co-cultures lined by human lung alveolar epithelium and microvascular endothelium interfaced across a rigid, porous, ECM-coated membrane and cultured in the same medium, we did not observe any of these classic hallmarks of RILI. Transwell cultures failed to mimic the hypertrophy of the alveolar cells (**Supplementary Fig. S2a,b**), increase in barrier permeability (**Supplementary Fig. S2c**) or changes in gene expression (**Fig. 2e**) observed in response to radiation exposure in the Alveolus Chips. At the RNA level (**Fig. 2e**), the endothelium in these static cultures overexpress the leukocyte adhesion marker ICAM-1, but did not display other gene expression changes that are characteristic of RILI in vivo which were observed on-chip (e.g., increased expression of HMOX-1, α-SMA, DDIT3, etc.). Similarly, while exposure of Transwell cultures to radiation induced an initial increase in cytokines like ICAM-1, MCP-1, IL-6 and IL-8 at 6 h post-radiation, the levels of these inflammatory cytokines returned to baseline levels by 24 h and there were no significant differences compared to controls (**Fig. 2f**). Furthermore, cells irradiated on Transwells exhibited a lower number of nuclear 53bp1 foci indicating reduced DNA damage in response to the same 16 Gy radiation exposure (**Fig. 2g**). Thus, these results demonstrate the importance of recreating the local physical microenvironment experienced by breathing lung alveolar tissues *in vivo,* including fluid flow and cyclic mechanical deformations (as well as an ALI), when attempting to model RILI *in vitro*.

Taken together, these studies established that the human breathing Lung Alveolus Chip can recapitulate key hallmarks of RILI *in vitro*, including DNA damage, junction disruption, increased barrier permeability and a heightened inflammatory response, whereas the same cells cultured under static conditions do not. Importantly, a radiation dose of 16 Gy and the presence of PBMCs were both necessary to capture this acute lung injury response on-chip.

### DNA damage and cell cycle arrest dominate early and are followed by inflammation

To further understand the mechanistic pathways underlying RILI in the human Lung Alveolus Chip, transcriptomic analyses were performed. At 6 h post radiation exposure, 217 genes were differentially expressed (21 upregulated and 127 downregulated) in chips exposed to 16 Gy versus controls (**Fig. 3a,b**). Interestingly, while 85 genes altered their expression in response to radiation exposure in both the epithelium and endothelium (**Supplementary Fig. S3a**), all but 3 of these common genes were upregulated rather than inhibited by this stimulus (**Supplementary Fig. S3b,c).** Unsupervised principal-component analysis (PCA) of RNA seq clustering data demonstrated two distinct gene expression clusters in both the epithelium and endothelium (**Supplementary Fig. S3d**). Gene set enrichment analysis (GSEA) of the hallmark pathways revealed downregulation of pathways involving the G2M checkpoint, E2F targets and mitotic spindle in the epithelium as well as endothelium (**Fig. 3c,d**), which is consistent with similar DNA damage to both lung tissues in response to radiation, as described above (**Fig. 1c,d**). However, the endothelium also exhibited upregulation of the immune response and allograft rejection pathways that generally represent a protective response to injury [18] (**Fig. 3d**). Gene ontology (GO) analysis similarly showed downregulation of cell division pathways relating to mitotic spindle formation, nuclear division, cytokinesis, and chromatid segregation in both lung tissue types (**Supplementary Fig. S3e**). However, we also observed a concomitant upregulation in the epithelial to mesenchymal transition (EMT) pathway that causes epithelial cells to lose their polarity and become more migratory and invasive into surrounding tissues [26].

**Fig. 3:**
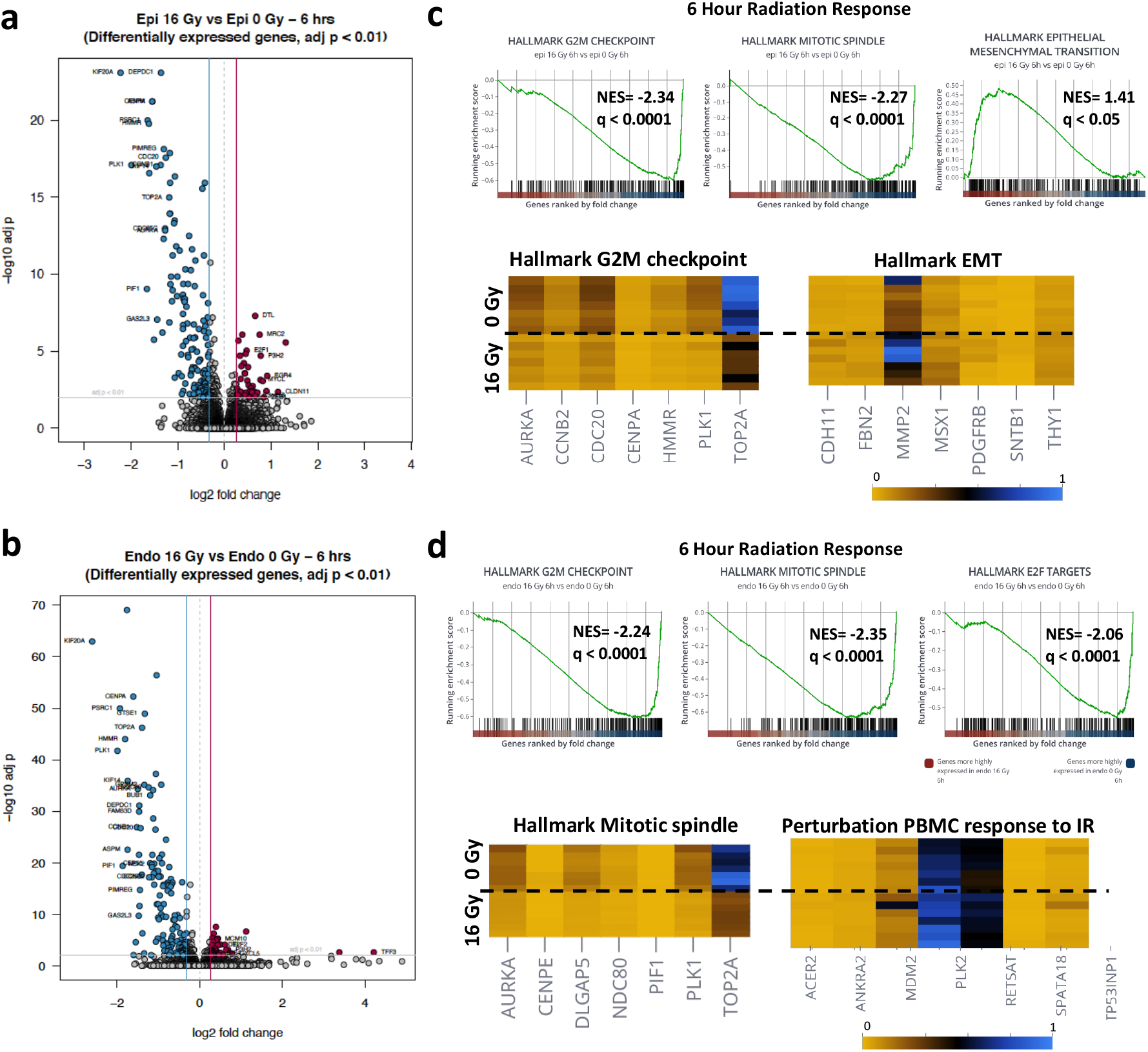
Transcriptomic analyses shows that DNA damage and cell cycle arrest are early effects of radiation. (a) Volcano plot of differentially expressed genes (DEGs) of alveolar epithelium, 6 h after radiation exposure (b) Volcano plot of differentially expressed genes (DEGs) of endothelium, 6 h after radiation exposure (c) Gene set enrichment analysis of the epithelium showing downregulation of hallmark G2M checkpoints, E2F targets and mitotic spindle, heatmaps showing top genes affected in the negative regulation of the G2M checkpoint and upregulation of EMT re (d) Gene set enrichment analysis of the endothelium showing downregulation of hallmark G2M checkpoints, E2F targets and mitotic spindle, heatmaps showing top genes affected in the negative regulation of the mitotic spindle organization and PBMC response to IR. Plots created with Pluto (https://pluto.bio).

In contrast, when we analyzed similar responses at 7 days after radiation exposure, greater changes in gene expression were observed in the vascular endothelium compared to the alveolar epithelium (**Fig. 4**). At this later time point, 595 genes were differentially expressed in the epithelium in response to radiation, whereas 1519 genes changed in the endothelium, with 111 genes changing in both (**Supplementary Fig. 4a)**. In contrast to the 6 hour time point, more genes were upregulated than downregulated in both tissue types (**Supplementary Fig. 4b,c**). Unsupervised PCA of RNA seq clustering data again revealed 2 distinct gene expression clusters in both tissue types in radiated Alveolus Chips compared to controls (**Fig. S4d,e**). Interestingly, GSEA demonstrated that the hallmark cell cycle pathways that were previously downregulated at 6 h were upregulated at 7 days (**Fig. S4f**). This is consistent with the analysis of reactome pathways that showed an increase in DNA repair, including base excision repair, and activation of ataxia telangiectasia and Rad3-related protein (ATR) in response to replication stress, as well as an increase in senescence and senescence-associated secretory phenotype. GSEA analysis of the KEGG pathways also revealed upregulation of cytosolic DNA sensing in response to radiation in the epithelium (**Fig. 4c**), most likely mediated by induction of ZBP1 [27]. Interferons are known to induce overexpression of ZBP1 [28], and they are upregulated in response to radiation in the epithelium as well (**Fig. 4a**). Cytokine-cytokine receptor interactions and chemokine signaling pathways are also upregulated at this late time after exposure to radiation and this was accompanied by increases in expression of several inflammatory markers, including IL-6, IL-8, interferon Lambda 1 (IFNL1), and C-C Motif Chemokine Receptor 1 (CCR1), as well as general upregulation of interferon pathways, in the alveolar epithelium (**Fig. 4a,c** and **Supplementary Fig. S4f**).

**Fig. 4:**
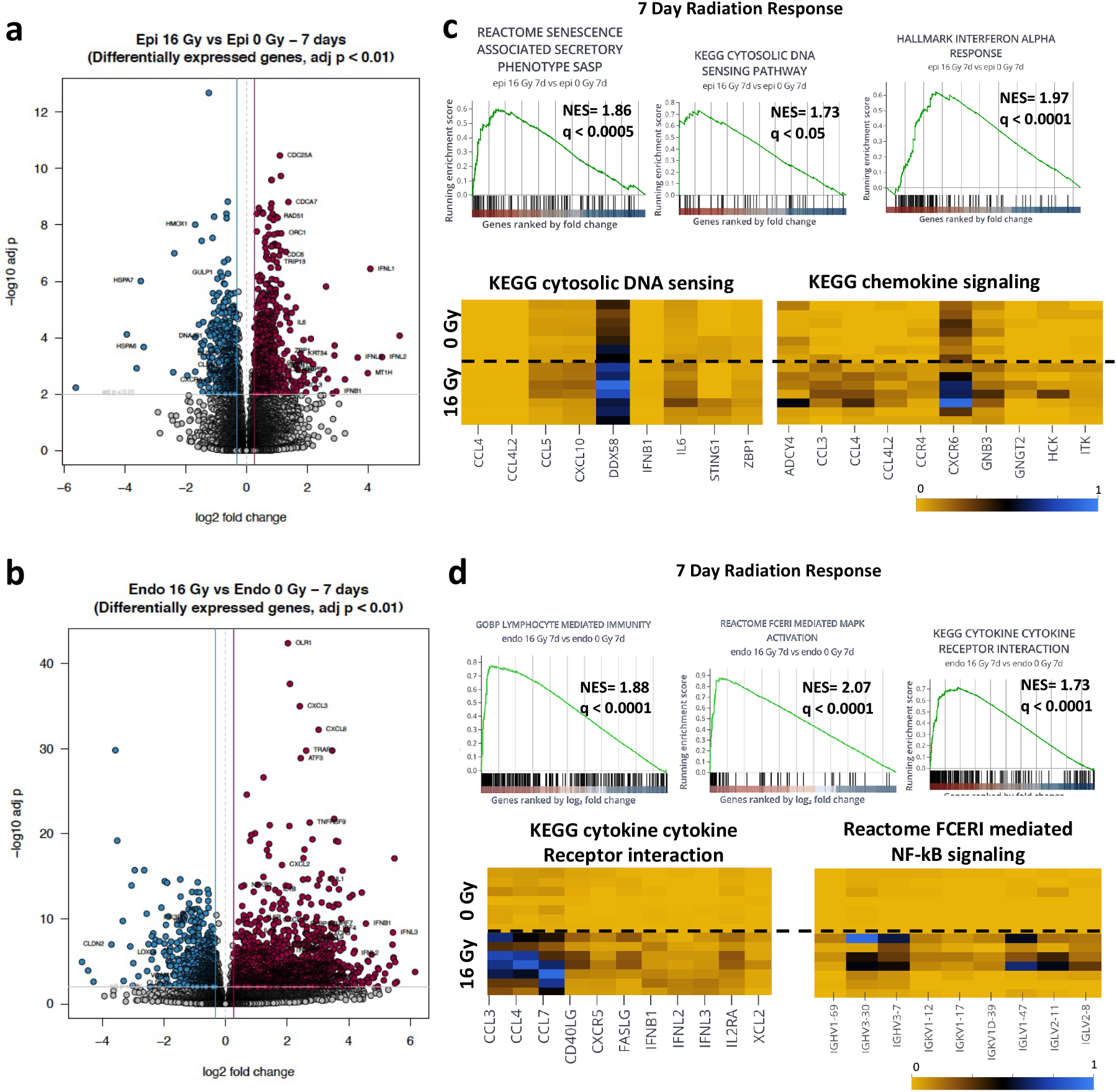
Progressive inflammation and higher endothelial susceptibility to radiation-injury 7d post radiation. (a) Volcano plot of differentially expressed genes (DEGs) of alveolar epithelium, 7 d after radiation exposure signaling (b) Volcano plot of differentially expressed genes (DEGs) of the endothelium 7 d after radiation exposure (c) Gene set enrichment analysis of the epithelium showing upregulation of hallmark interferon response, cytosolic DNA sensing pathways and senescence associated secretory pathways, heatmaps showing top genes affected in the upregulation of cytosolic DNA sensing and chemokine (d) Gene set enrichment analysis of the endothelium showing upregulation of lymphocyte mediated immunity, MAPK activation and increased cytokine-cytokine receptor interaction, heatmaps showing top genes affected in the upregulation of the cytokine expression and NF-κB signaling. Plots created with Pluto (https://pluto.bio).

In contrast, GSEA analysis of the vascular endothelium revealed significant upregulation of immune response, inflammatory response, and allograft rejection pathways mediated by TNF-α signaling via NF-κB, KRAs signaling, complement activation, IL2 STAT5 signaling, IL6 JAK-STAT3 and IFN-γ response pathways (**Fig. 4b** and **Supplementary Fig. S4f)**. Analysis of reactome pathways confirmed that the NF-κB pathway was upregulated in the endothelium within the irradiated Alveolus Chip, and that this was accompanied by progressive and sustained inflammation marked by the activation of late-stage inflammatory markers IL-1β, TGF-β, IL-10, and IL-16, together with an upregulation of CXCL8, CXCR3, and CXCL10 (**Fig. 4d** and **Supplementary Fig. S4e)**. We also observed increased expression of the potent vasoactive peptide endothelin-1 (EDN1) as well as oxidized low-density lipoprotein receptor-1 (OLR1), which is a key molecule associated with endothelial dysfunction during atherosclerosis progression. In addition, GO analysis revealed an increase in vasculature development pathways (**Supplementary Fig. S4f**), which is consistent with induction of a healing response to endothelial injury.

### HMOX-1 is a central regulator of radiation induced lung injury

Importantly, both transcriptomic analyses (**Fig. 3a,b**) and qPCR (**Fig. 5a**) revealed that HMOX-1, which converts heme to biliverdin, Fe2+, and CO as part of an anti-oxidant response (**Fig. 5b**)[29], is upregulated in both the epithelium and the endothelium at 6 h post radiation. Interestingly, however, while HMOX-1 levels dropped to baseline in the epithelium by day 7, they remained high in the endothelium at this time (**Fig. 5a**). Moreover, an independent analysis of the transcriptomics data using a Network Model for Causality-Aware Discovery (NeMoCAD) algorithm [30, 31] revealed that HMOX-1 was a central target in the top 10 (of 538) regulatory networks (**Supplementary Fig. S4g**). As the NeMoCAD algorithm has been previously used to predict therapeutic targets and repurpose drugs that reverse disease states[30, 31], this led to the hypothesis that the initial upregulation of HMOX-1 ensures radioprotection and that therapeutic elevation of HMOX-1 could alleviate radiation injury in the lung. But recent studies have indicated that HMOX-1 may play a dual role in lung injury and apart from its beneficial anti-oxidant and anti-inflammatory properties, as it also can promote cell death via ferroptosis [32]. Despite the upregulation of HMOX-1 at 6 h post radiation, ferroptosis markers were not differentially expressed in radiated samples (not shown). However, at 7 d post-irradiation, several ferroptosis markers are affected in the endothelium, and both downregulation of GPX4 and upregulation of PTGS2 and CHAC1 were observed (**Supplementary Fig. S5a**), indicating that ferroptotic pathways may be active at this time [33, 34].

**Fig. 5:**
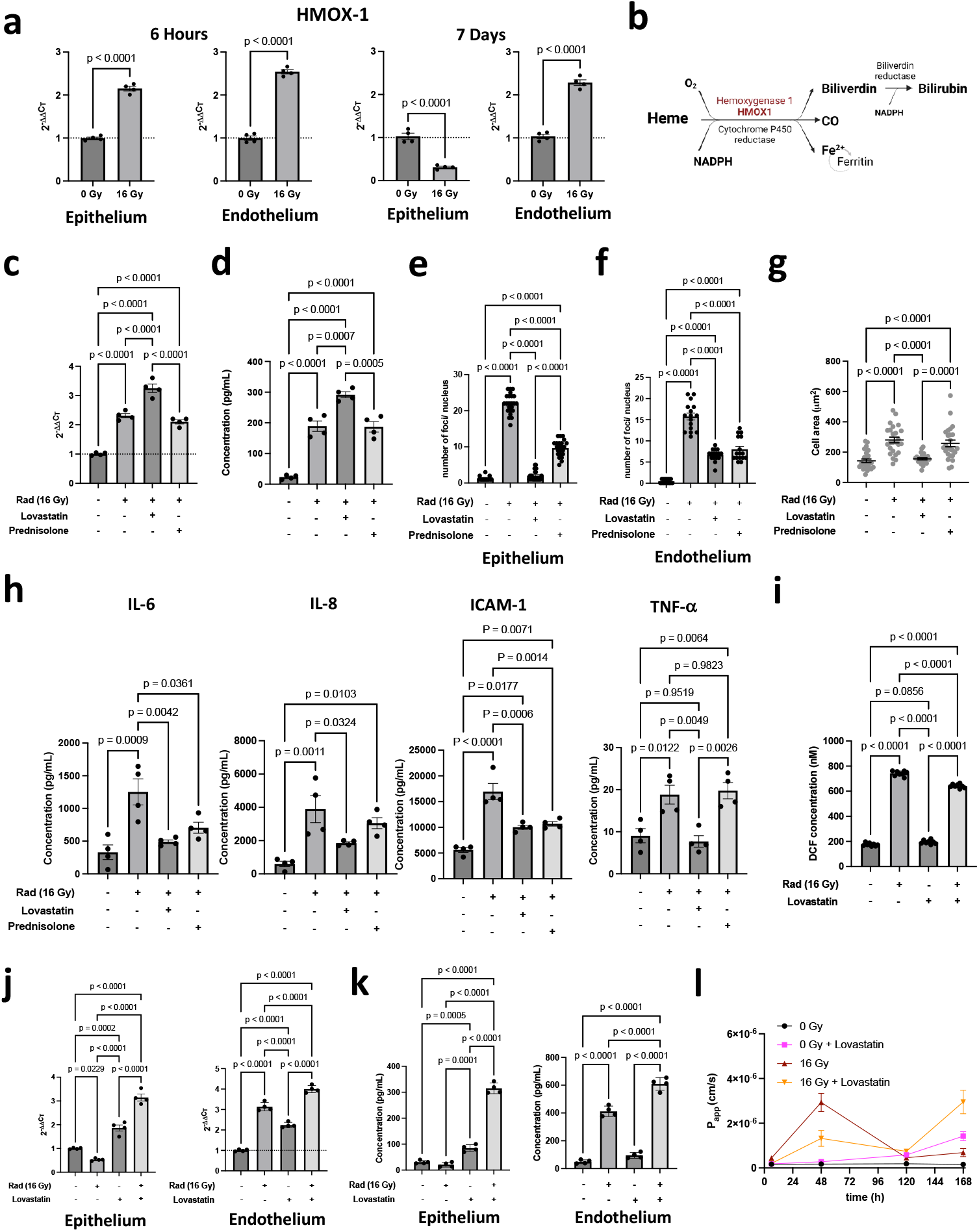
Identification of HO-1 as a therapeutic target for RILI and testing the effect of lovastatin and prednisolone on RILI. (a) gene expression levels of HMOX-1 in the epithelium and endothelium at 6 h and 7d post radiation, Two-tailed Student’s t-test with Welch’s correction (b) Pathway showing mechanism of action of HMOX-1 towards Bilirubin formation accompanied by the release of Fe2+ and CO (c) Effect of lovastatin and prednisolone of HMOX-1 RNA level, 6 h after radiation exposure and (d) on HO-1 protein, 24 h after radiation exposure. One-way ANOVA, n=4 chips (e) Quantification of 53BP1 foci/nucleus in the epithelium and (f) endothelium. (g) Effect of lovastatin and prednisolone on cellular hypertrophy in the alveolar epithelium (h) Effect of lovastatin and prednisolone on the cytokine response induced by radiation injury 24 h post-exposure. Representative panel showing the cytokines that were affected by the drug treatments. One-way ANOVA, n=4, DF=3, F(IL-6)= 10.7, F(IL-8)= 10.3, F(ICAM-1)=28.3 and F(TNF-α)= 12.0 (i) ROS assay 2 h after radiation exposure showing that presence of lovastatin decreases ROS levels in radiation exposed samples (j) HMOX-1 expression, 7 d post-radiation by qPCR and (k) HO-1 expression by ELISA assay, 7 post-radiation, indicating that HMOX-1 remains upregulated in both epithelium and endothelium in the presence of lovastatin (l) Changes in barrier permeability over time from 6 h to 7d post-radiation.

To directly investigate whether HMOX-1 plays an active role in RILI, we treated Lung Alveolus Chips with lovastatin 3h prior to radiation exposure because it has been reported to elevate HMOX-1 expression, particularly in the lung [35]. Indeed, lovastatin-treated chips showed higher levels HMOX-1 expression at both the gene (**Fig. 5c**) and protein (**Fig. 5d**) levels, and this was accompanied by significant suppression of acute radiation injury and cellular DNA damage, as indicated by a lower number of 53bp1 nuclear foci in both the epithelium and endothelium at 6 hours post-radiation (**Fig. 5e,f** and **Supplementary Fig. S5b**). Lovastatin also reduced epithelial cell hypertrophy (**Fig. 5g** and **Supplementary Fig. S5c**) and suppressed the inflammation response as indicated by a significant decrease in IL-6, IL-8, TNF-α, and ICAM-1 levels when measured 1 day after radiation exposure (**Fig. 5h**). As a positive control, we prophylactically treated chips with prednisolone prior to irradiation because it is commonly administered during radiotherapy to reduce inflammation. Indeed, prednisolone significantly reduced the number of 53bp1 foci both in the epithelium and endothelium (**Fig. 5e,f**). But while pretreatment with prednisolone decreased the levels of some cytokines (e.g., IL-6 and ICAM-1) at 24 h, IL-8 and TNF-α levels were still significantly higher than in control non-irradiated samples (**Fig. 5h**). Lovastatin treatment also reduced ROS levels compared to the no-drug radiation controls (**Fig. 5i**). Overall, the lovastatin treatment suppressed DNA damage and inflammation to a greater degree than prednisolone when measured during the first day after radiation exposure and it sustained this upregulation of HMOX-1 gene expression (**Fig. 5j**) and protein levels (**Fig. 5k**) for up to 7 d post radiation in both the epithelium and endothelium. Unfortunately, extended studies revealed that while lovastatin initially protected against tissue barrier disruption for at least up to 48 h after radiation injury, permeability increased significantly after 5 d (**Fig. 5l**) and this was accompanied by visible fluid accumulation in the epithelial channel (not shown). These disruptive effects of lovastatin occurred earlier than the barrier disruption observed in chips exposed to 16 Gy radiation in the absence of the drug (**Fig. 5l**).

## Discussion

Recent guidelines of the American Thoracic Society [36] suggest that animal models of acute lung injury induced by radiation should mimic 3 out of 4 criteria: 1) histopathologic evidence of tissue injury, 2) alteration of the alveolar-capillary barrier, 3) an inflammatory response, and 4) physiological dysfunction. Our results show that human Lung Alveolus Chips that experience dynamic fluid flow and breathing motions can be used to recapitulate these hallmark features of RILI observed in human lung when exposed to a radiation dose of 16 Gy *in vitro*. Specifically, we found that exposure to this high dose of radiation induces DNA damage, cellular hypertrophy, tight junction disruption, increased barrier permeability, ROS production, and subsequent fluid accumulation on-chip.

DNA damage is often characterized by analyzing DSB number and morphology, and we observed simpler DSBs at lower doses. This is consistent with the finding that as radiation dosage increases, the 53bp1 foci cluster into larger complexes that maybe repair centers [37]. We also observed induction of a senescence-associated secretory phenotype in response to radiation on-chip, which is another classic characteristic of radiation-pneumonitis in vivo [38]. Importantly, this Organ Chip model replicates progressive inflammation, as demonstrated by production of clinically relevant early (IL-6, IL-1α, TNF-α etc.) and late (IL-1 β, TGF-β, GM-CSF) stage cytokine markers of RILI, whereas this was not observed in a static Transwell culture. The finding that inflammation of the endothelium is also more prominent at day 7 after radiation exposure on-chip is interesting and is consistent with radiation-pneumonitis developing beginning ~1 week after exposure in radiated patients [2, 4].

Cytosolic DNA sensing has been previously reported to mediate radiation-induced lung injury when analyzing in vivo models and we also observed up regulation of genes associated with this pathway as well in the Alveolus Chip. This has been shown to be mediated by upregulation of ZBP1 [27], which is a Z-DNA-binding protein that may be induced by interferons [28], and ZBP1 also was upregulated in response to radiation in the Lung Chip model. In addition, we observed activation of inflammatory pathways, including TNF-α signaling via NF-κB, which was accompanied by FCERI mediated NF-kB signaling. Upregulation of EDN1 and OLR1 are also consistent with past studies that suggest endothelial apoptosis and ROS generation are upstream of radiation-induced epithelial cell injury and DNA damage responses [39, 40].

Finally, transcriptomic analysis combined with use of a machine learning-based computational network analysis algorithm (NemoCAD) [30, 31] suggested that HMOX-1 may play a central role in the RILI response, and hence, it could represent a potential therapeutic target. Additional experimental results confirmed that HMOX-1 is upregulated both in the epithelium and endothelium at 6 hours post-radiation, which is consistent with past work demonstrating that it is also upregulated and can be cytoprotective in other lung diseases (e.g., interstitial pneumonia, malaria-associated lung injury, silicosis) [41–43]. Importantly, when we used lovastatin to artificially upregulate HMOX-1 levels in Lung Alveolus Chips exposed to radiation, we observed significant suppression of DNA damage and cell hypertrophy as well as a decrease in inflammation at 24 h after exposure. Thus, these data suggest that HMOX-1 also plays a protective role in the lung’s initial response to radiation-induced injury. However, lovastatin treatment aggravated barrier disruption over time as indicated by disruption of the alveolar-capillary interface and increased fluid accumulation in the epithelial air channel.

Thus, while HMOX-1 may be critical for protection against acute radiation injury immediately following radiation exposure, prolonged elevation could be counterproductive. This may, in part, explain while lung injury only first manifests itself about 1 week after radiation exposure in vivo. Sustained upregulation of HMOX-1 in response to radiation exposure could lead to cellular accumulation of Fe^2+^ and thereby increase ferroptosis and inflammation. Ferroptosis inhibitors have been shown to provide protection against RILI in mice in vivo [44, 45]. This possibility is also supported by the findings that elevation of HMOX-1 appears to be associated with TGF-β-mediated lung inflammation [46] and that it can aggravate infection [32] and stimulate fibrosis [47, 48].

Taken together with our results, these observations suggest that HMOX-1 may serve dual roles during RILI: high levels of expression are required to protect against injury at early times, whereas continued elevation can exacerbate the injury response. Future radiation countermeasure strategies that target HMOX-1 must be designed to take the complexity of this dual response into account, for example, by using HMOX-1 agonists in the first few days after radiation exposure and antagonists at later times.

The human Lung Alveolus Chip RILI model offers many advantages over existing *in vitro* models because it more faithfully mimics key hallmarks of this disease. Perhaps most importantly, it replicates the sustained inflammation response that is central to RILI; however, it also has certain limitations. For example, past studies have shown that neutrophils and fibroblasts are important contributors to RILI and they are missing in our model. But due to the modular design of the chip, neutrophils, fibroblasts, and other potential cellular contributors can be integrated into the model in the future, as this has been done in the past [21–23, 49, 50]. Another limitation is that whole body radiation exposure induces injuries in multiple organs simultaneously (e.g., bone marrow, intestine) and there could be systemic cross talk that contributes to RILI from these distant sites via the circulation [51]. This is more easily studied in animal models, however, they do not faithfully mimic human organ radiation dose sensitivities or injury responses in the lung. Thus, future studies could be carried out in which the human Lung Alveolus Chip is linked fluidically with Organ Chip models via their vascular endothelium-lined channels as was done previously [51] to explore these systemic effects that are missing in the present study. Nevertheless, the results shown here demonstrate that many features of RILI observed in human patients can be replicated in vitro using the Alveolus Chip but not using more conventional static culture models. Thus, the Lung Alveolus Chip may represent a useful preclinical model for discovery of new radiation countermeasure drugs as well as for identification of potential biomarkers of RILI progression.

## ACKNOWLEDGEMENTS

This research was sponsored by funding from the US Food and Drug Administration grant 75F40119C10098, and the Wyss Institute for Biologically Inspired Engineering. The authors would like to thank B. Lubamba and S. Gilpin for their helpful advice during the early phase of this project and E. Javorsky for her editorial assistance.

## AUTHORSHIP CONTRIBUTIONS

### Contribution

Q.D. participated in the ideation, design and performance of all experiments and analyzed the data working with S.H. and D.E.I, who also supervised the work. A.J. participated in the performance of experiments, data collection and discussions. A.W. acquired preliminary data that formed the basis of early experiments. Q.D., R.M. and A.W. acquired confocal microscopy images. Q.D., S.H., and D.E.I. prepared the manuscript with input from all authors.

## METHODS

### Lung Alveolus cultures in microfluidic chips

Two-channel microfluidic chips (Chip-S1) and automatic fluid handling ZOE systems were obtained from Emulate Inc (Boston, MA, USA). The two channels in the chip device are separated by a porous (8 μm pore size), PDMS membrane. After activating the culture surface of the chip using ER1/ER2 following manufacturer’s instructions, the channels were coated with 200 μg/mL Collagen IV (5022-5MG, Advanced Biomatrix) and 15 μg/mL laminin (L4544-100 UL, Sigma) at 37 ^o^C overnight in a CO2 incubator (day 0). After coating is complete (day 1), chips are washed with DPBS (+ Ca^2+^, +Mg^2+^), lung microvascular endothelial cells (Lonza CC-2527, P5) are perfused at a seeding density of 8 × 10^6^ cells/mL onto the bottom channel of the chips, inverted and allowed to attach to the underside of the membrane for 1 h. Following this, alveolar epithelial cells (Cell Biologics H-6053, P1) are seeded onto the top channel of the chip at a seeding density of 1.6 × 10^6^ cells/mL and allowed to attach for 1 h. Following this, respective media are fed onto both the channels, and the chips are maintained under static conditions overnight. On day 2, the chips are inserted into Pods (fluid feeding chambers, Emulate Inc.) and connected to the ZOE systems to ensure perfusion. The apical and basal channels are perfused with Alveolar epithelial growth medium (Cell Biologics, H6621) and endothelial growth medium (Lonza, EGM2-MV, CC-3202), respectively at a flow rate of 45 μL/h. On day 5, the apical medium is supplemented with 1 μM dexamethasone to enhance barrier formation. Following this, on day 7, medium is removed from the apical channel to establish an ALI. Chips continued to be perfused with EGM2-MV in the basal channel. On day 9, the perfusing medium was changed to EGM2-MV with 0.5% FBS. On day 11, the parameters on the ZOE instrument were changed to apply cyclic 5% mechanical strain, at 0.25 Hz frequency to mimic breathing motions of the lung. The mechanical strain is applied for 4 days before the chips are ready for treatment. This 14-day incubation ensures an alveolar-capillary interface with good barrier function [22]. On day 15, a modified EGM2-MV (with 50 nM hydrocortisone, instead of 550 nM in the original formulation) is perfused into the basal channel prior to exposure to radiation.

### Exposure to radiation on chip

PBMCs (StemCell, #70025.1) were labeled with CytoTox green and perfused into the bottom channel at a density of 10^6^ cells/mL for 5 min at a flow rate of 500 mL/h. Following PBMC perfusion, chips and attached Pods were removed from the ZOE and exposed to gamma radiation using a Cesium source (Cs-137). The chips were exposed to radiation ranging from 12-16 Gy (**Fig. 1**). Experiments shown in **Fig. 2–5** include samples that were exposed to 16 Gy radiation. The sham irradiated control samples (0 Gy) were removed from the incubator and left at RT inside the biosafety cabinet. Following exposure to radiation, chips were connected back to the ZOE instrument and PBMCs were allowed to flow for 2 h at 500 μL/h. Following this, fresh medium was added to the bottom channel inlet and perfused at 45 μL/h.

### RNA extraction from alveolus Chip for qRT-PCR and RNA-seq

For RNA analyses, RNA was extracted at 6 h and 7 d post-radiation exposure to assess early and late effects of radiation, respectively. RNA isolation is performed using the RNeasy Plus Micro Kit (Qiagen 74034), following manufacturer’s instructions, except few modifications made to suit the alveolus chip system. Briefly, 10 μL of β-mercaptoethanol was added to 1 mL Buffer RLT plus. To extract RNA, an empty 200 μL filtered barrier tip was inserted at the top channel outlet and washed with 100 μL DPBS (+ Ca^2+^, +Mg^2+^) to collect apical washes. After collecting the apical channel washes, another empty 200 μL filtered tip was inserted into the apical channel outlet. The basal channel outlet and inlet are plugged with empty 200 μL filtered tips. The tops of these two tips are plugged with the help of two fingers and 200 μl of Buffer RLT plus BME was introduced through the apical channel inlet, using a 200 μL filtered tip. The lysate was homogenized by pipetting the liquid 3-5 times through the apical channel. The same technique was repeated for the basal channel containing the endothelium. Lysates from each channel were collected in RNase free tubes (1.5 mL) and stored at −80 °C for RNA-seq analysis, or qPCR. RNA isolation was performed following manufacturer’s instructions after this step.

### RT-qPCR

Total RNA was isolated using the RNeasy Micro Plus Kit (Qiagen 74034) for isolation from chips or RNeasy Mini Plus Kit (Qiagen 74106) for isolating RNA from transwells or well-plates. RNA concentrations were quantified using a nanodrop spectrophotometer (Thermo Scientific^™^ 912A1100). After quantification, 100-300 ng of RNA was used for cDNA synthesis. Reverse transcription was achieved using iScript cDNA Synthesis kit (Bio-Rad 1708891). Following this, quantitative real time PCR (qRT-PCR) was conducted using TaqMan^™^ Fast Advanced Master Mix (Thermo Fisher 444455) and analyzed on Quantstudio 7 Flex Real-Time PCR system (Thermo Fisher 4485701), following manufacturer’s instructions with UNG incubation hold at 50 °C for 2 min ➔ Polymerase activation Hold at 95 °C for 2 min ➔ PCR (40 cycles) ➔ Denature at 95 °C for 1 min and anneal/extend at 60 °C for 20 s. Primer-specificity was confirmed by melting curve analysis. Relative RNA levels were quantified by the ΔΔC_t_ method [52] and normalized to endogenous controls of HPRT1 (for lung alveolar epithelium),and B2M or GAPDH (for microvascular endothelium). All Taqman probes were purchased from Thermo Fisher Scientific (Supplementary Information **Table S1**).

### Immunostaining and confocal microscopy

At the designated end-points, typically 6 h and 7d after radiation exposure (unless specified otherwise), cells were fixed with 1 % paraformaldehyde (PFA) in warm medium and incubated at RT for 5 min. Subsequently, they were fixed with 4% PFA (in PBS(+/+)) for an additional 30 min at RT. Followed by washing with PBS (-/-) and stored at 4 °C or processed immediately for imaging. For immunostaining, the cells were permeabilized with 0.1% Triton-X-100 in PBS for 10 min, and blocked with 5% donkey serum or 1% BSA/PBS for 1 h at RT. The cells are then incubated with primary antibodies at specific dilutions, overnight at 4 °C under shaking conditions. Fluorescent probe-conjugated secondary antibodies were incubated (in case of unconjugated primary antibodies) with the cells, followed by nuclear staining with Hoechst. The list of primary antibodies and their respective dilutions are listed in Supplementary **Table S2**. All immunofluorescence images are representative and have been captured in at least 2 separate experiments.

### Reactive oxygen species (ROS) assay

To test ROS levels in the alveolus chip in response to radiation, the OxiSelect^™^ In vitro ROS/RNS assay (catalog no. STA-347) was used, following manufacturer’s instructions. Briefly the assay employs a quenched fluorescent probe dichlorodihydrofluorescin DiOxyQ (DCFH-DiOxyQ). The probe is primed by a quench removal agent and stabilized to the DCFH form. ROS and RNS can react with DCFH, and gets rapidly oxidized to the fluorescent 2’,7’-dichlorodihydrofluorescein (DCF). Fluorescence intensity of DCF is directly proportional to the ROS/RNS levels in the sample. Standard curve of known DCF concentration vs fluorescence intensity is plotted and used to calculate DCF concentration from the sample effluents.

### Permeability assay

To test permeability of the epithelial-endothelial barrier of the chips, tracer dye molecule, FITC-dextran (40 kDa) was perfused to basal inlet reservoir of the pods, at a concentration of 100 μg/mL of each tracer. The apical inlet reservoir was filled with fresh medium. The flow rate of the apical and basal compartments were set to 120 μL/h. After 2 h of perfusion, effluents were collected from the apical and basal outlets and concentrations of the tracers quantified by fluorescence spectroscopy on a BioTek well-plate reader. Apparent permeability (Papp) calculations were based on chip manufacturer’s (Emulate) instructions.

### Cytokine analysis

Effluents from the basal (vascular) outlets were collected 24 h and 7 d after radiation exposure, and were analyzed for a 13-panel of cytokines, and chemokines like IL-1α, IL-6, IL-β, IL-1β, GM-CSF, TNF-α, MCP-1, MIP-1α, IP-10, ICAM-1, P-selectin, Serpine1, using customized ProcartaPlex assay kits (Invitrogen).The analyte concentrations are evaluated using a Bio-Plex 3D suspension array system and analyzed for standard curve fitting and concentration calculations with Bio-Plex Manager software (Bio-Rad, v 6.0). TGF-β (Cat. no. DB100B) and HMOX-1 (Cat no. OKBB00836) were measured using ELISA kits (R & D Systems).

### Bulk RNA sequencing

RNA-seq was performed by Azenta Life Sciences using a RNA-seq package including polyA selection and sequencing using an Illumina HiSeq for 150 bp pair-ended reads. These sequence reads were trimmed using Trimmomatic v.0.36 to eliminate adapter sequences and nucleotides with poor quality. Subsequently, these trimmed reads were mapped onto the Homo sapiens GRCh38 reference genome using STAR aligner v.2.5.2b. Unique gene hit counts were computed using Counts from the Subread package v.1.5.2 followed by differential expression analysis. Significant DEGs were defined as log_2_ (fold change) ≥ 0.5 and p < 0.01 to adjust for false discovery. Differential expression analysis was performed by comparing the different groups. Genes were filtered to include genes with at least 3 reads in at least 20% of samples in any group. Differential expression analysis was then performed using the DESeq2 R package [53], which tests for differential expression based on a model using negative binomial distribution. Log2(fold change) was calculated for each comparison. Thus, genes with a positive log2(fold change) value had increased expression, while genes with a negative log2(fold change) value had decreased expression in the radiated (16 Gy) samples. Volcano plots showing the log2(fold change) of each gene on the x-axis and the −log10(adjusted p-value) on the y-axis. Points are filled (in blue or red) if the gene’s adjusted p-value is ≤ 0.01. The false discovery rate (FDR) method was applied for multiple testing correction [54]. FDR-adjusted p-values are shown on the y-axis of the volcano plot. An adjusted p-value of 0.01 was used as the threshold for statistical significance. Analysis and figures for transcriptomic analyses were generated using Pluto (https://pluto.bio).

### NemoCAD analysis

NemoCAD is a machine-learning based computational tool to analyze transcriptomics signatures that can be targeted to reverse the global transcriptomic changes observed due to radiation exposure [30, 31]. This approach is agnostic because it is run without an *a priori* defined drug/ gene target or mechanism of action. NemoCAD uses previously computed drug-gene or gene-gene interaction probabilities and DEG signatures of the treated or diseased states and corresponding controls. The algorithm identifies target compounds that can revert a transcriptional signature in the diseased state to one observed in the healthy control.

### Statistics

All reported experiments were performed at least 3 times, with at least 4 technical repeats in each group. Data is represented as mean ±S.D. unless specified in the figure legend. Graph plotting and statistics were performed on GraphPad Prism (Version 9.4.0) Statistical significance for 2 group comparisons were determined using two-tailed Student’s t-test, using Welch’s correction for unpaired t-test. For multiple comparisons, one way ANOVA was used with Dunnett’s multiple comparison test when compared to a single control group. All n, p-values and F, df values are mentioned in the respective figure or the associated legends.

## REFERENCES

1. Society, A., Cancer treatment & survivorship facts & figures 2019–2021. 2019, American Cancer Society Atlanta.

2. Arroyo-Hernández, M., et al., Radiation-induced lung injury: current evidence. BMC pulmonary medicine, 2021. 21(1): p. 1–12.

3. Giuranno, L., et al., Radiation-induced lung injury (RILI). Frontiers in oncology, 2019. 9: p. 877–892.

4. Bledsoe, T.J., S.K. Nath, and R.H. Decker, Radiation pneumonitis. Clinics in chest medicine, 2017. 38(2): p. 201–208.

5. Magana, E. and R.E. Crowell, Radiation pneumonitis successfully treated with inhaled corticosteroids.(Case Report). Southern medical journal, 2003. 96(5): p. 521–525.

6. Abratt, R.P., et al., Pulmonary complications of radiation therapy. Clinics in chest medicine, 2004. 25(1): p. 167–177.

7. Beach, T.A., et al., Modeling radiation-induced lung injury: lessons learned from whole thorax irradiation. International journal of radiation biology, 2020. 96(1): p. 129–144.

8. Huang, W., et al., Acute proteomic changes in lung after WTLI in a mouse model: identification of potential initiating events for delayed effects of acute radiation exposure. Health physics, 2019. 116(4): p. 503–515.

9. Jackson, I.L., et al., A preclinical rodent model of radiation induced lung injury for medical countermeasure screening in accordance with the FDA animal rule. Health physics, 2012. 103(4): p. 463–473.

10. Rogers, C.J., et al., Identification of miRNA signatures associated with radiation-induced late lung injury in mice. PloS one, 2020. 15(5): p. e0232411.

11. Dabjan, M.B., et al., A survey of changing trends in modelling radiation lung injury in mice: bringing out the good, the bad, and the uncertain. Laboratory Investigation, 2016. 96(9): p. 936–949.

12. MacVittie, T.J., et al., Acute radiation-induced lung injury in the non-human primate: A review and comparison of mortality and co-morbidities using models of partial-body irradiation with marginal bone marrow sparing and whole thorax lung irradiation. Health Physics, 2020. 119(5): p. 559–587.

13. MacVittie, T.J., et al., The time course of radiation-induced lung injury in a nonhuman primate model of partial-body irradiation with minimal bone marrow sparing: clinical and radiographic evidence and the effect of neupogen administration. Health Physics, 2019. 116(3): p. 366–382.

14. Cohen, J., Vaccine studies stymied by shortage of animals. Science, 2000. 287(5455): p. 959–960.

15. Thakur, P., et al., Clinicopathologic and transcriptomic analysis of radiation-induced lung injury in nonhuman primates. International Journal of Radiation Oncology* Biology* Physics, 2021. 111(1): p. 249–259.

16. Araya, J., et al., Ionizing radiation enhances matrix metalloproteinase-2 production in human lung epithelial cells. American Journal of Physiology-Lung Cellular and Molecular Physiology, 2001. 280(1): p. L30–L38.

17. Jung, J.-W., et al., Ionising radiation induces changes associated with epithelial-mesenchymal transdifferentiation and increased cell motility of A549 lung epithelial cells. European journal of cancer, 2007. 43(7): p. 1214–1224.

18. Oshi, M., et al., Association of allograft rejection response score with biological cancer aggressiveness and with better survival in triple-negative breast cancer (TNBC). Journal of Clinical Oncology, 2021. 39(15_suppl): p. 561–561.

19. Jain, A., et al., Primary human lung alveolus-on-a-chip model of intravascular thrombosis for assessment of therapeutics. Clinical pharmacology & therapeutics, 2018. 103(2): p. 332–340.

20. Plebani, R., et al., Modeling pulmonary cystic fibrosis in a human lung airway-on-a-chip. Journal of Cystic Fibrosis, 2021. 21(4): p. 606–615.

21. Hassell, B.A., et al., Human organ chip models recapitulate orthotopic lung cancer growth, therapeutic responses, and tumor dormancy in vitro. Cell reports, 2017. 21(2): p. 508–516.

22. Bai, H., et al., Mechanical control of innate immune responses against viral infection revealed in a human lung alveolus chip. Nature communications, 2022. 13(1): p. 1–17.

23. Si, L., et al., Clinically relevant influenza virus evolution reconstituted in a human lung airway-on-a-chip. Microbiology Spectrum, 2021. 9(2): p. e00257–21.

24. Shibata, A. and P.A. Jeggo, Roles for 53BP1 in the repair of radiation-induced DNA double strand breaks. DNA repair, 2020. 93: p. 102915.

25. Costes, S.V., et al., Spatiotemporal characterization of ionizing radiation induced DNA damage foci and their relation to chromatin organization. Mutation Research/Reviews in Mutation Research, 2010. 704(1-3): p. 78–87.

26. Zhou, Y.-C., et al., Ionizing radiation promotes migration and invasion of cancer cells through transforming growth factor-beta–mediated epithelial–mesenchymal transition. International Journal of Radiation Oncology* Biology* Physics, 2011. 81(5): p. 1530–1537.

27. Takaoka, A., et al., DAI (DLM-1/ZBP1) is a cytosolic DNA sensor and an activator of innate immune response. Nature, 2007. 448(7152): p. 501–505.

28. Fu, Y., et al., Cloning of DLM-1, a novel gene that is up-regulated in activated macrophages, using RNA differential display. Gene, 1999. 240(1): p. 157–163.

29. Campbell, N.K., H.K. Fitzgerald, and A. Dunne, Regulation of inflammation by the antioxidant haem oxygenase 1. Nature Reviews Immunology, 2021. 21(7): p. 411–425.

30. Novak, R., et al., Target-agnostic discovery of Rett Syndrome therapeutics by coupling computational network analysis and CRISPR-enabled in vivo disease modeling. bioRxiv, 2022: p. 2022–03.

31. Sperry, M.M., et al., Target-agnostic drug prediction integrated with medical record analysis uncovers differential associations of statins with increased survival in COVID-19 patients. medRxiv, 2022: p. 2022–04.

32. Yang, S., et al., A Dual Role of Heme Oxygenase-1 in Tuberculosis. Frontiers in Immunology, 2022. 13: p. 1–12.

33. Chen, X., et al., Characteristics and biomarkers of ferroptosis. Frontiers in cell and developmental biology, 2021. 9: p. 637162.

34. Stockwell, B.R., Ferroptosis turns 10: Emerging mechanisms, physiological functions, and therapeutic applications. Cell, 2022. 185(14): p. 2401–2421.

35. Hsu, M., et al., Tissue-specific effects of statins on the expression of heme oxygenase-1 in vivo. Biochemical and biophysical research communications, 2006. 343(3): p. 738–744.

36. Kulkarni, H.S., et al., Update on the features and measurements of experimental acute lung injury in animals: An official American Thoracic Society workshop report. American Journal of Respiratory Cell and Molecular Biology, 2022. 66(2): p. e1–e14.

37. Neumaier, T., et al., Evidence for formation of DNA repair centers and dose-response nonlinearity in human cells. Proceedings of the National Academy of Sciences, 2012. 109(2): p. 443–448.

38. Ungvari, Z., et al., Ionizing radiation promotes the acquisition of a senescence-associated secretory phenotype and impairs angiogenic capacity in cerebromicrovascular endothelial cells: role of increased DNA damage and decreased DNA repair capacity in microvascular radiosensitivity. Journals of Gerontology Series A: Biomedical Sciences and Medical Sciences, 2013. 68(12): p. 1443–1457.

39. Dong, F., et al., Endothelin-1 enhances oxidative stress, cell proliferation and reduces apoptosis in human umbilical vein endothelial cells: role of ETB receptor, NADPH oxidase and caveolin-1. British journal of pharmacology, 2005. 145(3): p. 323–333.

40. Wei, J., et al., The role of NLRP3 inflammasome activation in radiation damage. Biomedicine & Pharmacotherapy, 2019. 118: p. 109217.

41. Pereira, M.L., C.R. Marinho, and S. Epiphanio, Could heme oxygenase-1 be a new target for therapeutic intervention in malaria-associated acute lung injury/acute respiratory distress syndrome? Frontiers in Cellular and Infection Microbiology, 2018. 8: p. 161–179.

42. Nakashima, K., et al., Regulatory role of heme oxygenase-1 in silica-induced lung injury. Respiratory research, 2018. 19(1): p. 1–11.

43. Sato, T., et al., Heme oxygenase-1, a potential biomarker of chronic silicosis, attenuates silica-induced lung injury. American journal of respiratory and critical care medicine, 2006. 174(8): p. 906–914.

44. Li, X., X. Zhuang, and T. Qiao, Role of ferroptosis in the process of acute radiation-induced lung injury in mice. Biochemical and Biophysical Research Communications, 2019. 519(2): p. 240–245.

45. Li, X., et al., Ferroptosis inhibitor alleviates Radiation-induced lung fibrosis (RILF) via down-regulation of TGF-β1. Journal of inflammation, 2019. 16: p. 1–10.

46. Ye, Q., et al., Heme Oxygenase 1 Contributes to TGF-β1-Induced Lung Myofibroblast Differentiation, in A107. MECHANISMS OF FIBROSIS/FIBROBLAST BIOLOGY. 2022, American Thoracic Society. p. A2324–A2324.

47. Hara, Y., et al., Heme oxygenase-1 in patients with interstitial lung disease: A review of the clinical evidence. The American Journal of the Medical Sciences, 2021. 362(2): p. 122–129.

48. Atzori, L., et al., Attenuation of bleomycin induced pulmonary fibrosis in mice using the heme oxygenase inhibitor Zn-deuteroporphyrin IX-2, 4-bisethylene glycol. Thorax, 2004. 59(3): p. 217–223.

49. Huh, D., G.A. Hamilton, and D.E. Ingber, From 3D cell culture to organs-on-chips. Trends in cell biology, 2011. 21(12): p. 745–754.

50. Huh, D., et al., Reconstituting organ-level lung functions on a chip. Science, 2010. 328(5986): p. 1662–1668.

51. Novak, R., et al., A robotic platform for fluidically-linked human body-on-chips experimentation. Nature biomedical engineering, 2020. 4(4): p. 407–420.

52. Livak, K.J. and T.D. Schmittgen, Analysis of relative gene expression data using real-time quantitative PCR and the 2-ΔΔCT method. methods, 2001. 25(4): p. 402–408.

53. Love, M.I., W. Huber, and S. Anders, Moderated estimation of fold change and dispersion for RNA-seq data with DESeq2. Genome biology, 2014. 15(12): p. 1–21.

54. Benjamini, Y. and Y. Hochberg, Controlling the false discovery rate: a practical and powerful approach to multiple testing. Journal of the Royal statistical society: series B (Methodological), 1995. 57(1): p. 289–300.

