## Supplementary Information for "Hemoxygenase-1 as a key mediator of acute radiation pneumonitis revealed in a human lung alveolus-on-a-chip"

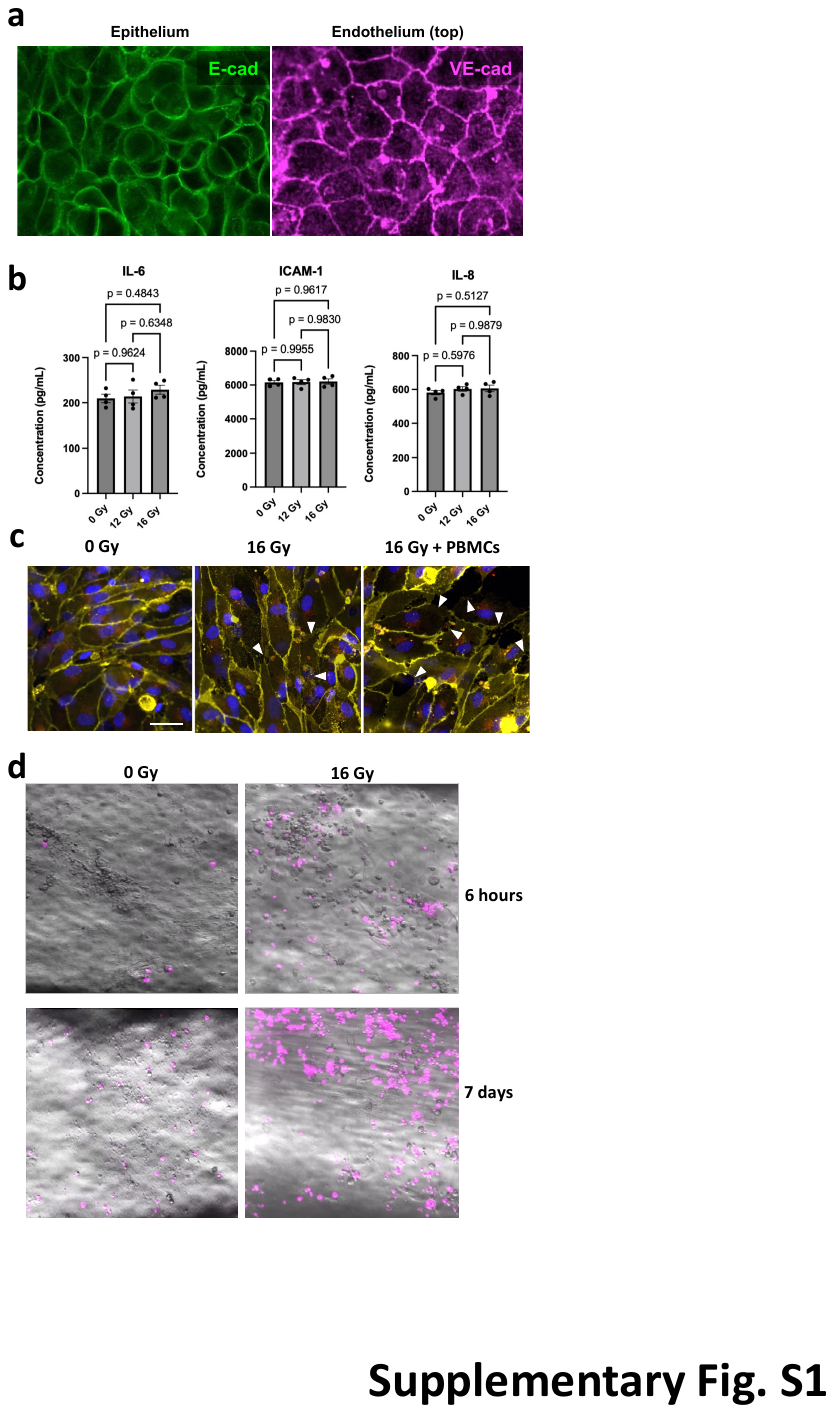

**Fig. S1:** (a) Confocal images of epithelial cells stained for E-cadherin(green) and endothelial cells stained for VE-cadherin adherens junction protein (magenta). Scale bar = 20 μm. (b) Representative Comparison of cytokine levels, 24 h post-radiation showed that 12 Gy and16 Gy radiation did not show an elevation in cytokine levels, in the absence of PBMCs (c) Presence of PBMCs in the endothelial compartment during radiation aggravates tight junction disruption (d) Fluorescence microscopy images showing PBMC recruitment to the endothelial side at 6 h and 7d post-radiation.

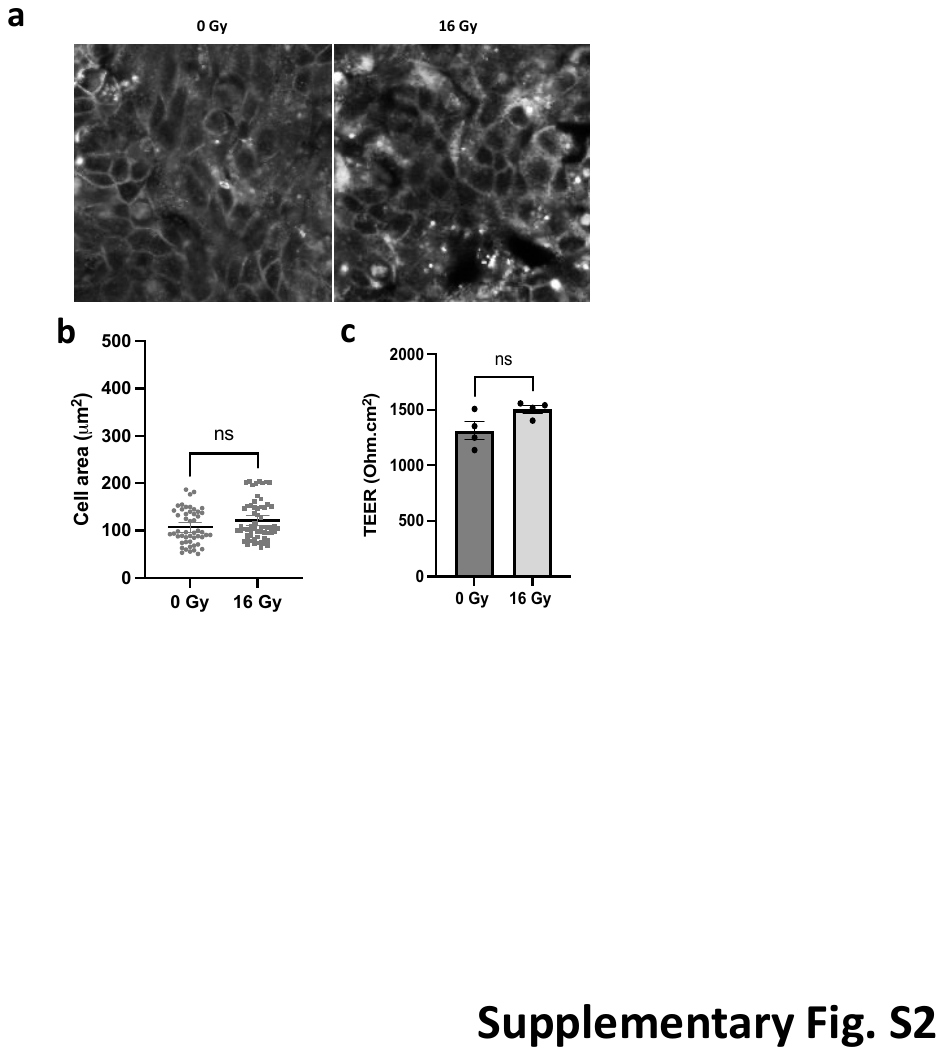

**Fig. S2:** (a) Representative Immunofluorescence images of alveolar epithelium in Transwell (E-cad staining). Alveolar cells on transwells do not exhibit hypertrophy in response to 16 Gy radiation (b) Quantification of cell size from E-cad staining images (data plotted from 8 different frames in each condition, n=98 for 0 Gy and n=78 for 16 Gy) (c) Assessment of barrier function shows no difference in the TEER resistance values in response to radiation on transwell.

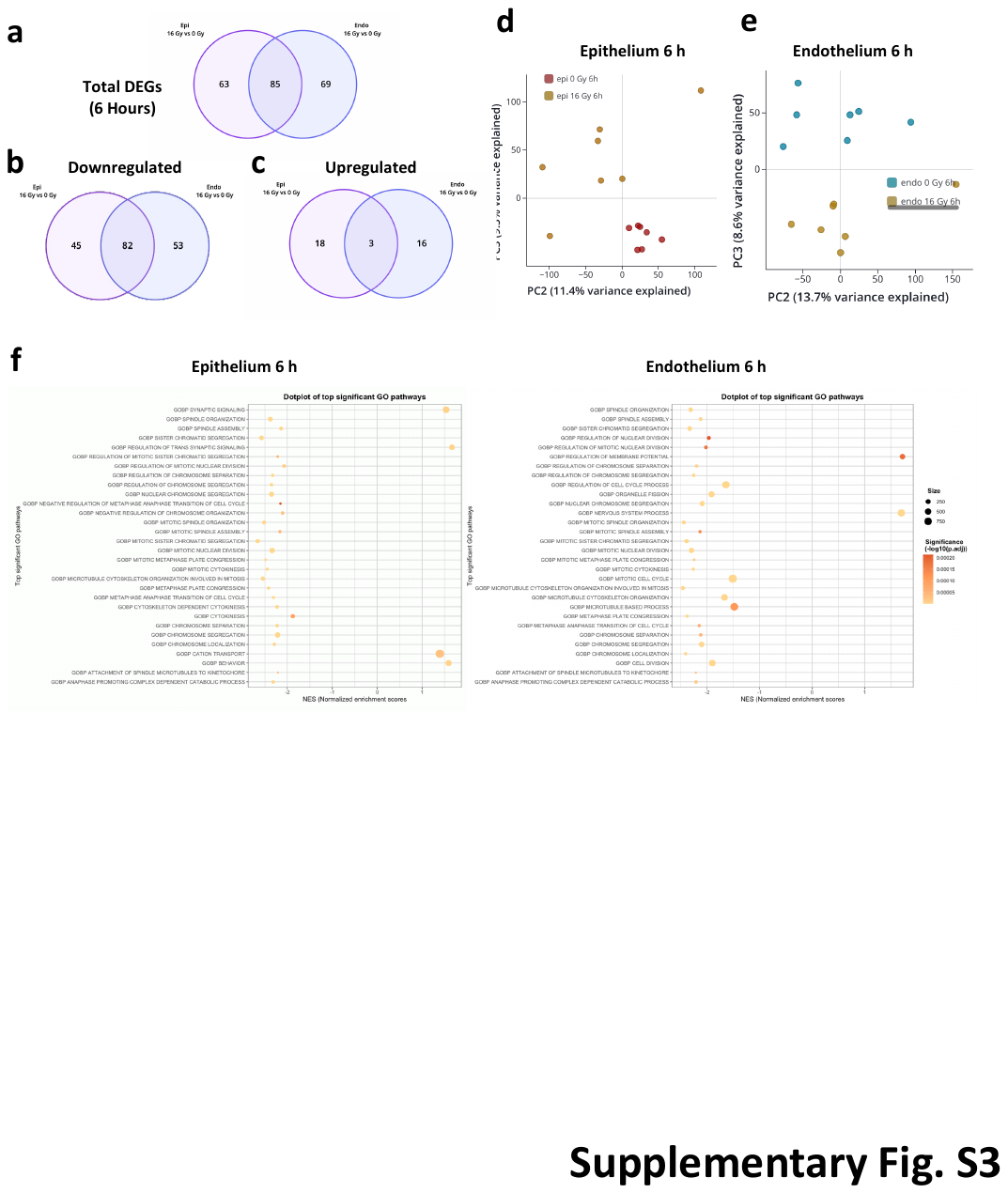

**Fig. S3:** Transcriptomic analyses at 6 h post-radiation. (a) Venn diagram showing total differentially expressed genes (DEGs) in response to radiation, 6 h after exposure, (b) Genes downregulated in the epithelium and endothelium, 6 h after radiation (c) Genes upregulated in the epithelium and endothelium, 6 h after radiation. Principal component analysis of the (d) epithelium and (e) endothelium showing the different clusters formed in response to radiation injury. (f) Dotplots showing the top significant gene ontology: biological pathways upregulated in response to radiation in the epithelium and endothelium

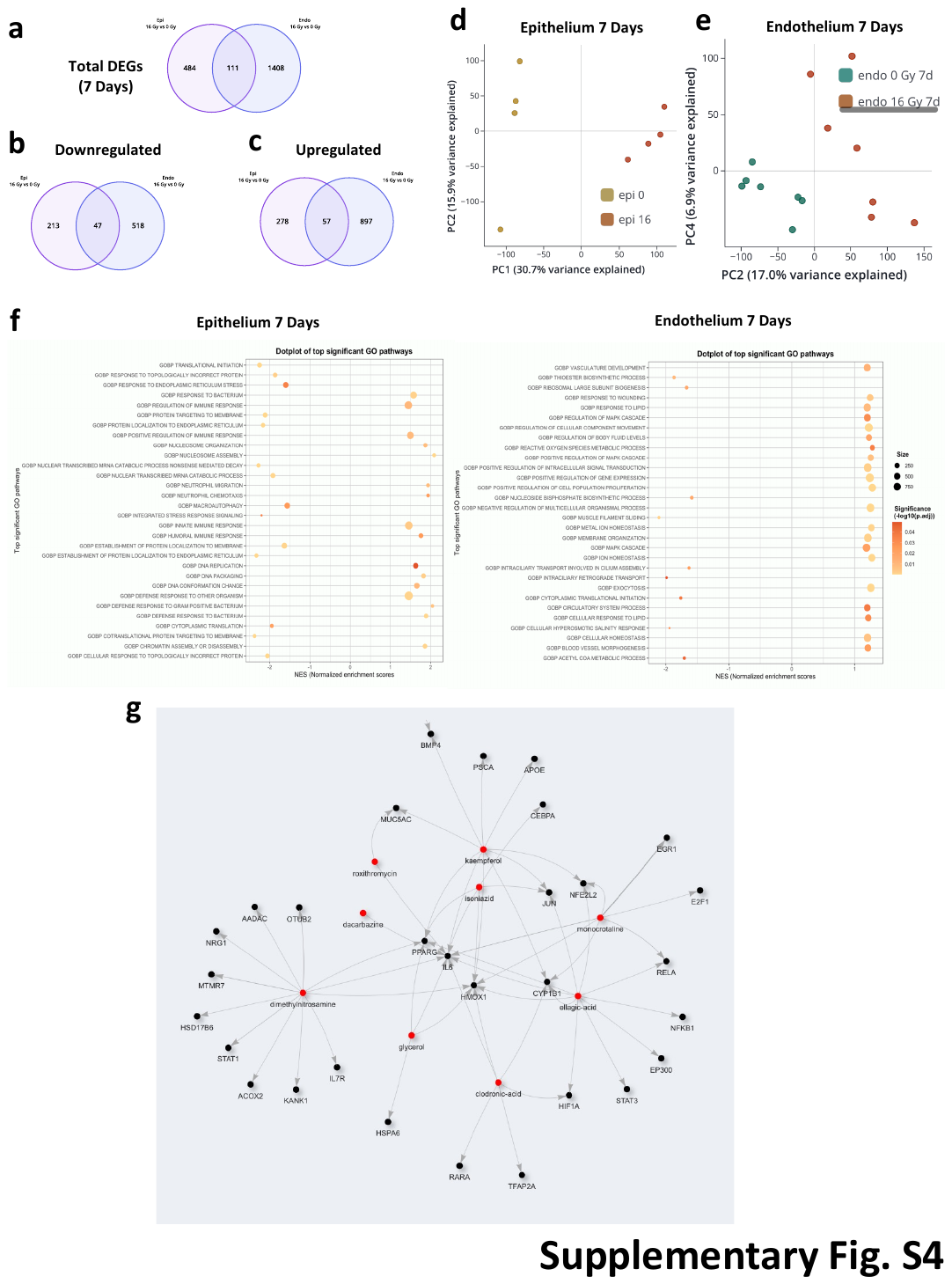

**Fig. S4**: Transcriptomic analyses at 7d post-radiation. (a) Venn diagram showing total differentially expressed genes (DEGs) in response to radiation, 7 d after exposure, (b) Genes downregulated in the epithelium and endothelium, 7 d after radiation (c) Genes upregulated in the epithelium and endothelium, 7 d after radiation Principal component analysis of the (d) epithelium and (e) endothelium showing the different clusters formed in response to radiation injury. (f) Dotplots showing the top significant gene ontology: biological pathways upregulated in response to radiation in the epithelium and endothelium (g) NeMoCAD analysis of the transcriptomic analyses at 7d post radiation showing that HMOX-1 and IL6 are identified as central targets in an agnostic approach.

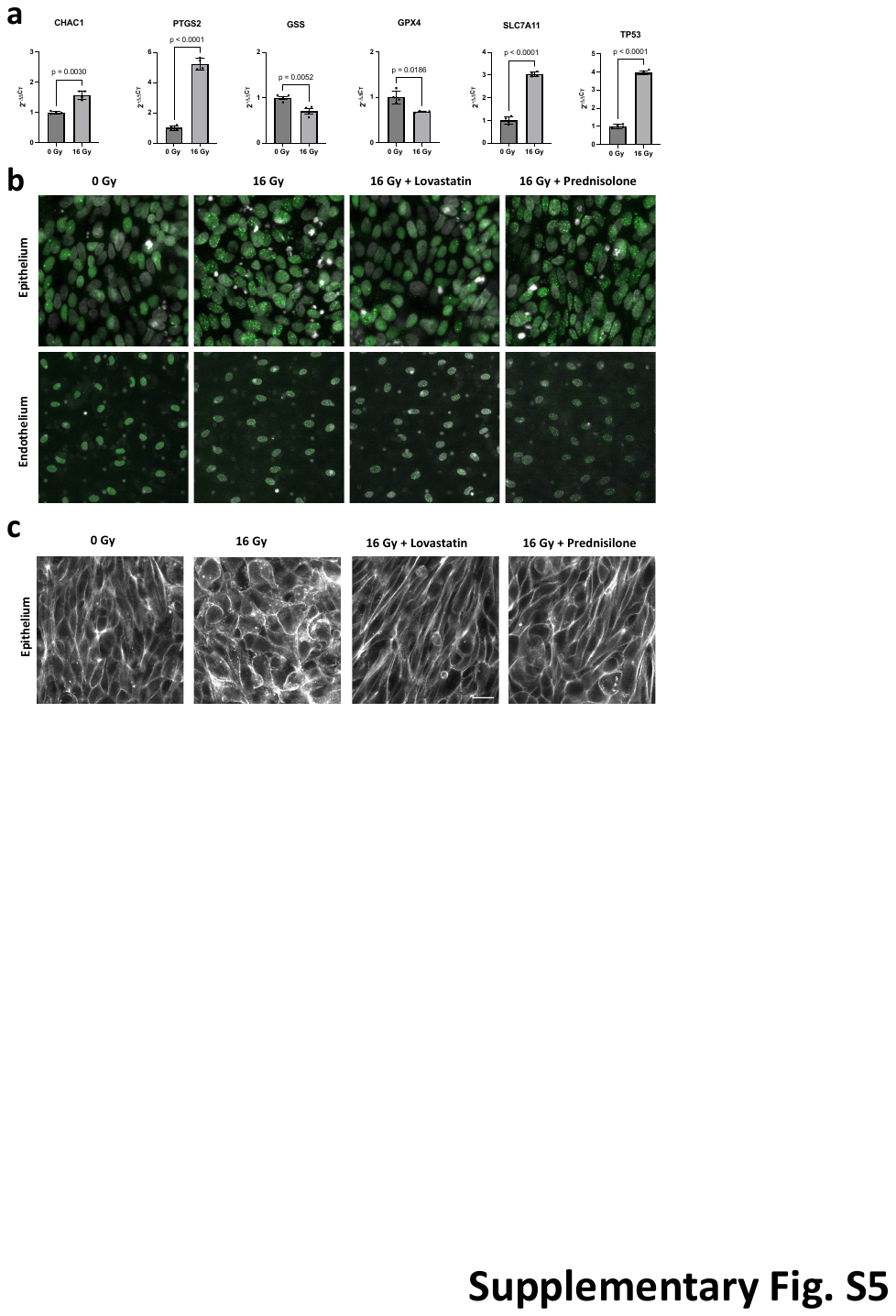

**Fig. S5:** (a) RNA expression of ferroptosis-associated markers in response to radiation at 7 d post-exposure. (b) IF imaging showing the effect of lovastatin and prednisolone on formation of 53bp1 nuclear foci and (c) cellular hypertrophy observed in the epithelium.

**Table S1. List of Thermo Fisher Taqman probes**

| **Gene symbol** | **Cat. no.** | **Assay ID** |
| --- | --- | --- |
| HPRT1 | 4331182 | Hs02800695_m1 |
| SNAI1 | 4331182 | Hs00195591_m1 |
| B2M | 4351370 | Hs00187842_m1 |
| AQP5 | 4331182 | Hs00387048_m1 |
| SFTPC | 4331182 | Hs00161628_m1 |
| SFTPB | 4331182 | Hs00167036_m1 |
| ACTA2 | 4331182 | Hs00909449_m1 |
| PDPN | 4331182 | Hs00366766_m1 |
| ICAM1 | 4331182 | Hs00164932_m1 |
| SELE | 4331182 | Hs00950401_m1 |
| ATM | 4331182 | Hs00175892_m1 |
| GPX4 | 4331182 | Hs00989766_g1 |
| H2AFX | 4331182 | Hs00266783_s1 |
| DDIT3 | 4331182 | Hs00358796_g1 |
| CHAC1 | 4331182 | Hs00225520_m1 |
| PTGS2 | 4331182 | Hs00153133_m1 |
| HMOX1 | 4331182 | Hs01110250_m1 |
| P21 | 4331182 | Hs01371942_m1 |
| TP53 | 4331182 | Hs01034249_m1 |
| BAX | 4331182 | Hs00180269_m1 |
| BCL2 | 4331182 | Hs04986394_s1 |
| GSS | 4331182 | Hs00609285_m1 |
| SLC7A11 | 4351372 | Hs00921933_m1 |

**Table S2. List of antibodies used in immunostaining**

| **Antibody** | **Company** | **Cat. No.** |
| --- | --- | --- |
| Anti-E Cadherin antibody | Abcam | ab1416 |
|  | Novus Biologicals | FAB748R-025 |
| Anti-ZO1 tight junction protein antibody - C-terminal | Abcam | ab190085 |
| Anti-CD31 antibody [JC/70A] | Abcam | ab9498 |
| Anti-53BP1 antibody | Abcam | ab36823 |
|  | Novus Biologicals | NB100-305AF594 |
| VE-Cadherin | Cell Signaling Technologies | 2500S |
|  | Novus Biologicals | FAB9381X |
